# Genomic, Ancestral and Networking Analyses of a High-Altitude Native American Ecuadorian Patient with Congenital Insensitivity to Pain with Anhidrosis

**DOI:** 10.1101/529263

**Authors:** Andrés López-Cortés, Ana Karina Zambrano, Patricia Guevara-Ramírez, Byron Albuja Echeverría, Santiago Guerrero, Eliana Cabascango, Andy Pérez-Villa, Isaac Armendáriz-Castillo, Jennyfer García-Cárdenas, Verónica Yumiceba, Gabriela Pérez-M, Paola E. Leone, César Paz-y-Miño

## Abstract

Congenital insensitivity to pain with anhidrosis (CIPA) is an extremely rare autosomal recessive disorder characterized by insensitivity to pain, inability to sweat and intellectual disability. CIPA is caused by mutations in the neurotrophic tyrosine kinase receptor type 1 gene (NTRK1) that encodes the high-affinity receptor of nerve growth factor (NGF). Patients with CIPA lack the primary afferents and sympathetic postganglionic neurons leading to lack of pain sensation and the presence of anhidrosis, respectively. Herein, we conducted a genomic analysis of 4,811 genes and 18,933 variants, including 54 mutations of NTRK1 in a high-altitude indigenous Ecuadorian patient with CIPA. As results, the patient presented 87.8% of Native American ancestry, 6.6% of African ancestry and 5.6% of European ancestry. The mutational analysis of the kinase domain of NTRK1 showed two pathogenic mutations, rs80356677 (Asp674Tyr) and rs763758904 (Arg602*). The genomic analysis showed 68 pathogenic and/or likely pathogenic variants in 45 genes, and two variants of uncertain significance in CACNA2D1 (rs370103843) and TRPC4 (rs80164537) genes involved in the pain matrix. The GO enrichment analysis showed 28 genes with relevant mutations involved in several biological processes, cellular components and molecular functions. In addition, the protein-protein interaction (PPi) networking analysis showed that NTRK1, SPTBN2 and GRM6 interact with several proteins of the pain matrix. In conclusion, this is the first time that a study associates genomic, ancestral and networking data in a high-altitude Native American Ecuadorian patient with consanguinity background in order to better understand CIPA pathogenesis.

## INTRODUCTION

Congenital insensitivity to pain with anhidrosis (CIPA), also known as hereditary sensory and autonomic neuropathy Type IV (HSAN-IV) (OMIM #256800), is an extremely rare autosomal recessive disorder that belongs to the HSAN family which are clinically and genetically heterogeneous disorders characterized by axonal atrophy affecting the sensory and autonomic neurons(1,2). HSAN-IV is characterized by recurrent episodes of unexplained fever, absence of reaction to noxious (or painful) stimuli, self-mutilating behavior, complete anhidrosis, variable degrees of intellectual disability(3), humoral immunodeficiency(4), palmoplantar keratoderma(5,6), and early onset renal disease(7). This condition occurs with an incidence of 1 in 125 million newborns(8,9).

Brain regions with pain perception are complex and have been best described as a pain matrix(10,11). According to Foulkes *et al* (2008), it consists of four phases in which different genes/proteins are involved(11). Nerves inside the skin have the ability to transmit the sensation of heat, cold and mechanical stimulation; Na^+^ and K^+^ channels drive nerve stimuli; the synaptic transmission occurs in the spinal cord via neurotransmitter receptors and Ca^2+^ channels; lastly, central, peripheral and microglia modulation occurs in brain(11). Nevertheless, patients with CIPA may present genetic alterations causing functional disruption in one of these pain matrix phases.

Only some hundred of cases of CIPA have been reported worldwide(8,12). The first reference to a similar pathology was mentioned by Dearborn in 1932(13), and it was published in 1963 by Swanson(14). Tunçbilek *et al* (2005) determined three clinical representative findings: insensitivity to pain, inability to sweat and intellectual disability(15). Indo *et al* (1996) associated CIPA pathogenesis with genetic loss-of-function mutations of the NTRK1 (neurotrophic receptor tyrosine kinase 1) gene(16). Grills and Schuijers (1998) postulated that nerve growth factor (NGF) function disruption also causes an altered process of fracture consolidation(17). Indo *et al* (2001) determined that CIPA is not only an autosomal recessive disorder, but also a uniparental disomy(18). Jarade *et al* (2002) observed ocular manifestations(19). Many studies of Weier *et al* (1995), Miura *et al* (2000), Indo *et al* (2001), Mardy *et al* (2001), Bonkowsky *et al* (2003) and Lin *et al* (2010) discovered novel mutations and polymorphisms in NTRK1 causing CIPA(20–24). Schreiber *et al* (2005) analyzed insulin-related difficulties(25). Brandes and Stuth (2006) evaluated anesthetic considerations(26). Many studies of Tanaka *et al* (1990), Indo (2002) and Melamed *et al* (2004) determined that NGF receptor failure causes a deficient development of dorsal root neurons (pain and temperature sensory system) autonomic sympathetic neural system (eccrine sweat glands innervation)(27–29). Abdulla *et al* (2014) observed heterotopic ossification and callus formation following fractures and eventually Charcot’s joint(30). Franco *et al* (2016) proposed that mutations of NTRK1 generate different levels of cell toxicity, which may provide an explanation of the variable intellectual disability observed in CIPA(31). Finally, Altassan *et al* (2016) identified novel NTRK1 mutations in CIPA individuals through exome DNA sequencing(1).

NTRK1, also known as TRKA, is located on chromosome 1q21-22 and encodes the neurotrophic tyrosine kinase-1 receptor, which is autophosphorylated in response to the NGF, thus, activating various pathways of intracellular signaling transduction such as cell growth, differentiation and survival(1,27,32). These signal transduction pathways mediate innervation of the skin by sensory and sympathetic axons(33). Additionally, anhidrosis is explained by the lack of the sympathetic postganglionic neurons, and the pain insensitivity is caused by the absence of the NGF-dependent primary afferents(1,34). According to the Human Gene Mutation Database and the ClinVar, NTRK1 has ~79 alterations among single nucleotide polymorphisms (SNPs), insertions and deletions, inherited in an autosomal recessive pattern(35,36). Indo *et al* (1996) has reported for the first time NTRK1 mutations associated with CIPA in an Ecuadorian family(37). However, this is the first time that clinical, ancestral, genomic, PPi networking and gene ontology (GO) enrichment analyses were performed in a high-altitude Native American (indigenous) patient with CIPA disease and family consanguinity background.

### Case presentation

The case of a 9-year-old girl who was born in the community of Piaba (2,418 meters above sea level, MASL), in Cotacachi, located in the north of Ecuador, is presented. She is diagnosed with CIPA, which begins to be suspected after 72 hours of life, where she developed fever of unknown origin; consequently, she was admitted to the hospital, she stayed there for 26 days and she was discharged without specific diagnosis. After one month of age, she presented recurring episodes of fever.

When she was 4 months old, she was diagnosed with pneumonia; while she stayed at the hospital, her neurodevelopment was examined by means of the Denver test which provided the following result: unusual for her age. The neurological examination showed generalized hypotonia, active symmetrical movements, incomplete cephalic support, absence of pain sensitivity during peripheral line placement, normal deep tendon reflex and absence of corneal reflex. Additionally, it was observed that there was a 1-centimeter dermal ulcer, with regular edges on the proximal and distal phalange of the first finger of the right hand, and lesions in healing process on the second finger; consequently, with general anesthesia, a skin biopsy of the sternal region was carried out. The results showed superficial and deep dermis with some cutaneous appendices constituted by hair follicles and sweat glands that in multiple cuts have no innervation zones, which can be corroborated since the mother presents absence of perspiration. The immunohistochemical staining test for nerve fibers with S100 protein was negative (Fig 1a).

**Fig 1.**
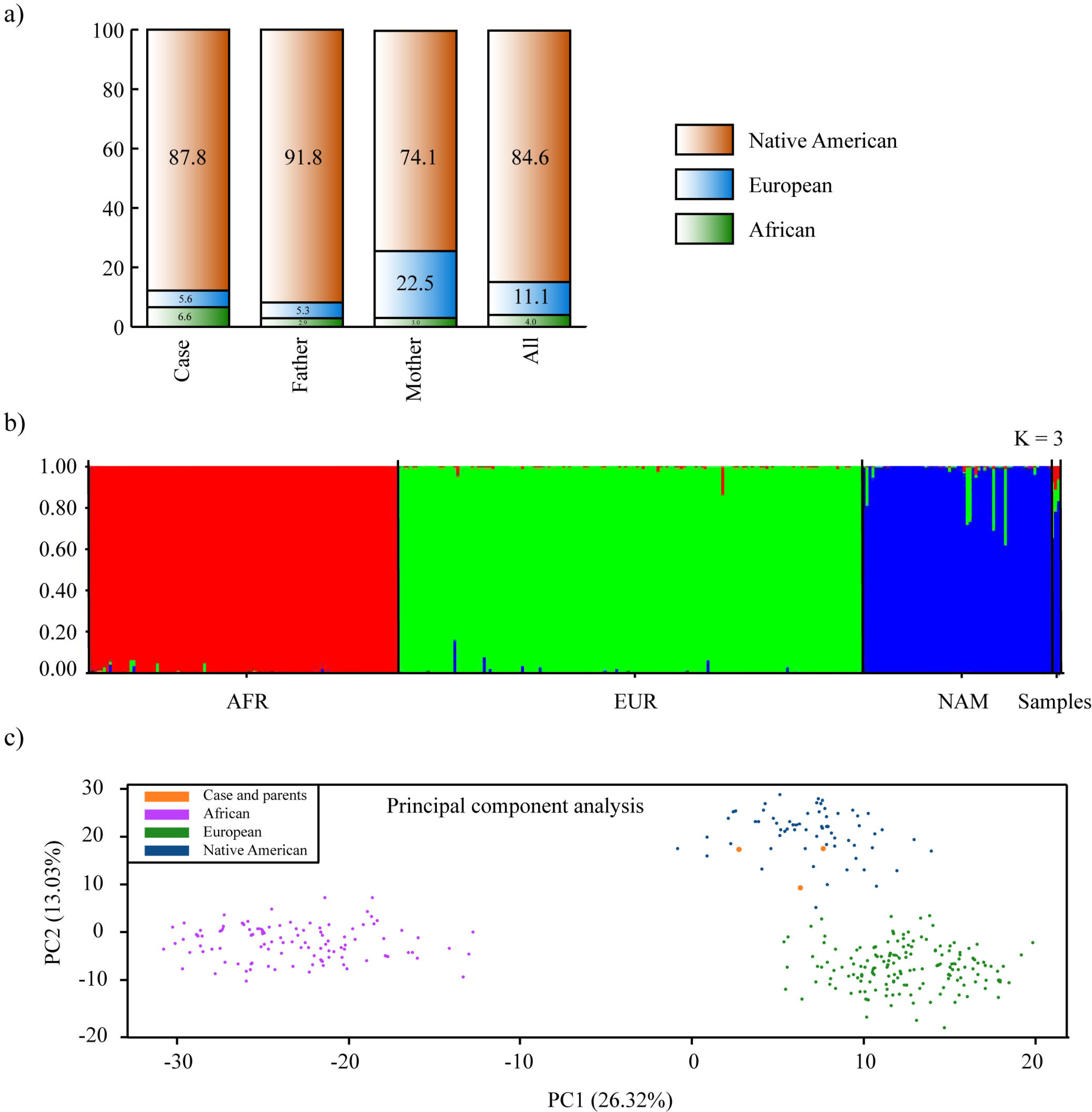
Clinical features of CIPA patient. a) Skin biopsy shows a thin epidermis with hyperkeratosis, there are few sebaceous glands and no nerve terminals are observed. b) Several fractures in tibia and femur. c) Karyotype. d) Family genealogical tree.

When she was 16 months old, she was diagnosed with Riga-Fede disease because she presented ulcerative plaques in the oral mucosa and tongue deformities. Furthermore, due to previous ulcers, the patient developed osteomyelitis on the distal phalanges of first and second fingers of the left hand (positive culture for *Staphylococcus aureus* sensitive to cephalexin). At 2.5 years old, and later at 3 years and 2 months old she presented bilateral corneal ulcers of traumatic origin. At 6 years and 4 months old she suffered a tibia fracture caused by falling (Fig 1b). At 6 years and 7 months she presented a distal tibial fracture without a determined cause. During the consolidation process a mass was found in the fracture area; a biopsy was carried out and osteochondroma was diagnosed. At 8 years and 1 month she broke her left femur because of a fall. Finally, at 8 years and 5 months old she suffered a subtrochanteric fracture of the right femur, requiring surgery (Fig 1b).

Additionally, a cytogenetic test was performed, resulting in normal karyotype 46, XX (Fig 1c).

As for the family background, a sister of the patient passed away at 18 months old after developing fever of unknown origin. Fig 1d details the genealogical tree, and consanguinity between relatives of both parents.

## RESULTS

### Clinical data

Patient shows various injuries such as dermal ulcer on the proximal and distal phalange. Additionally, she was diagnosed with Riga-Fede disease because she presented ulcerative plaques in the oral mucosa and tongue deformities. Fig 1a shows a skin biopsy of the sternal region performed under general anesthesia. The sample showed superficial and deep dermis with some cutaneous appendices constituted by hair follicles and sweat glands, which, in multiple cuts, did not have attached innervation zones. The immunohistochemical staining for S100 protein was negative. This test confirmed the diagnosis of the patient. Fig 1b shows a sequence of several fractures in the tibia and femur, reaching several complications due to its late treatment due to the absence of pain. Fig 1c shows the cytogenetic analysis. 25 metaphases with 46 chromosomes were scored obtaining a normal karyotype 46, XX. Fig 1d shows the genealogical tree of the patient with CIPA and her family, in which there is the presence of consanguinity among family members, and a sister who died due to constant fever but without an accurate diagnosis.

### Ancestry informative markers

The admixture analysis comparing the family under study with the reference population gave us a result that Native American ancestry is the most prevalent with 84,6%, followed by European ancestry 11,1% and lastly African ancestry with 4% (Fig 2a and 2b). On the other hand, the principal component analysis displayed in the first two components 39.35% of the variance. There is a clear separation with Africans and Europeans, being the family members into the Native American cluster (Fig 2c).

**Fig 2.**
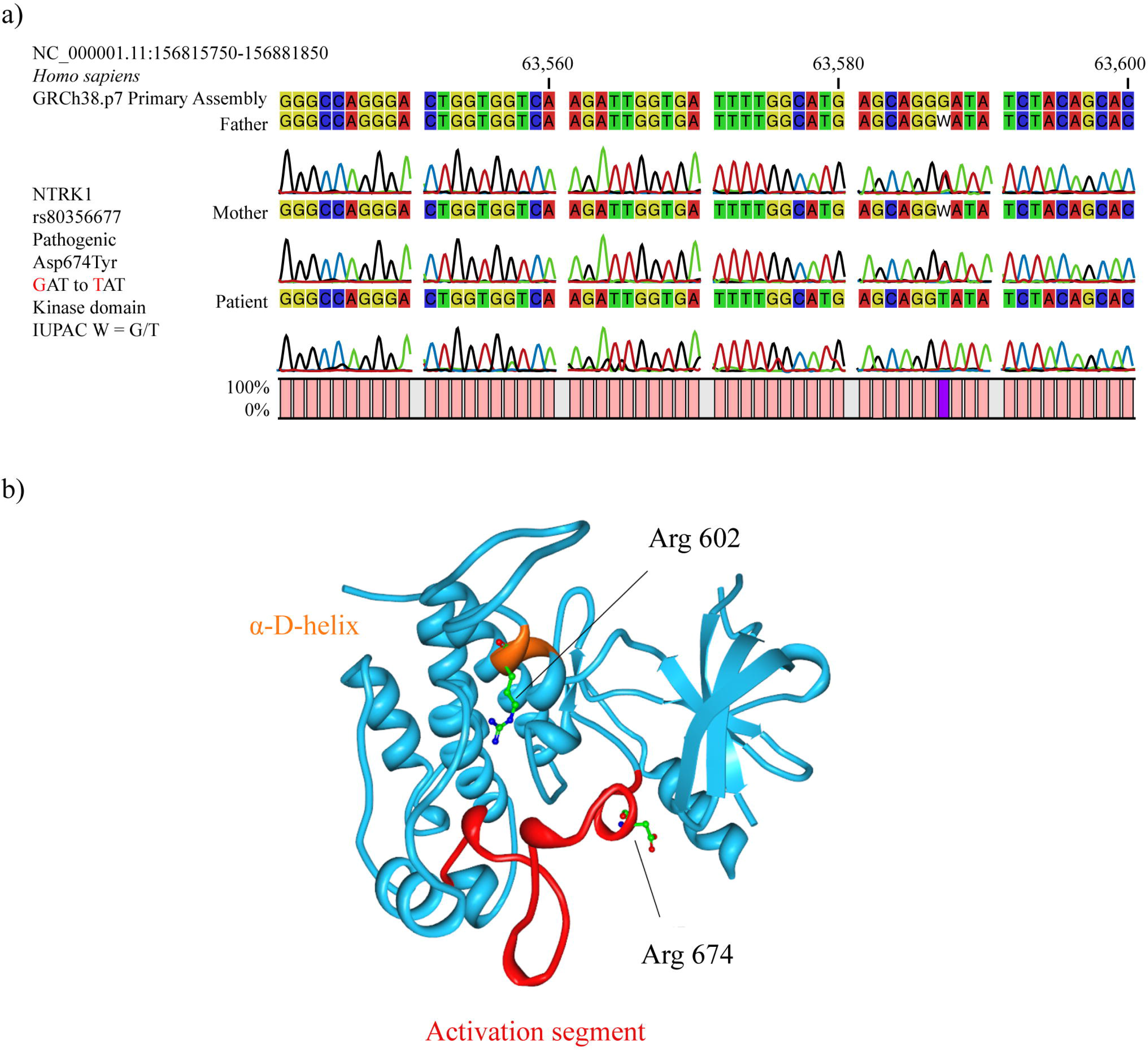
Analysis of ancestry informative markers. a) Percentage of ancestry per each family member. b) Bar plot of ancestry obtained from reference populations and Ecuadorian samples (EUR, European; AFR, African; NAM, Native American; ECU, Ecuadorians). Results from Structure v.2.3.4 with K = 3 and assuming migration model. c) Principal component analysis of Ecuadorian samples and reference populations. Plot constructed using RStudio.

### Mutational analysis of NTRK1

After carrying out the sequence analysis (Sanger) of the 14 regions that comprise the 54 analyzed mutations of the NTRK1 gene of the CIPA patient and her parents, the patient was observed to have the homozygous mutant genotype T/T of the missense pathogenic mutation rs80356677 (Arg674Tyr) in the kinase domain of NTRK1. At the amino acid level, this mutation changes the aspartic acid (Asp) by tyrosine (Tyr) at position 674, while at the nucleotide level there is a change of the guanine (G) to thymine (T), being the GAT codon changed by TAT. Regarding parents, both have the heterozygous genotype G/T of the pathogenic mutation rs80356677. That is, there is a pattern of autosome recessive inheritance (Fig 3a). In regard to the protein structure of NTRK1, the rs80356677 (Arg674Tyr) mutation is located in the activation segment, which is phosphorylated in protein kinases to activate its function (Fig 3b).

**Fig 3.**
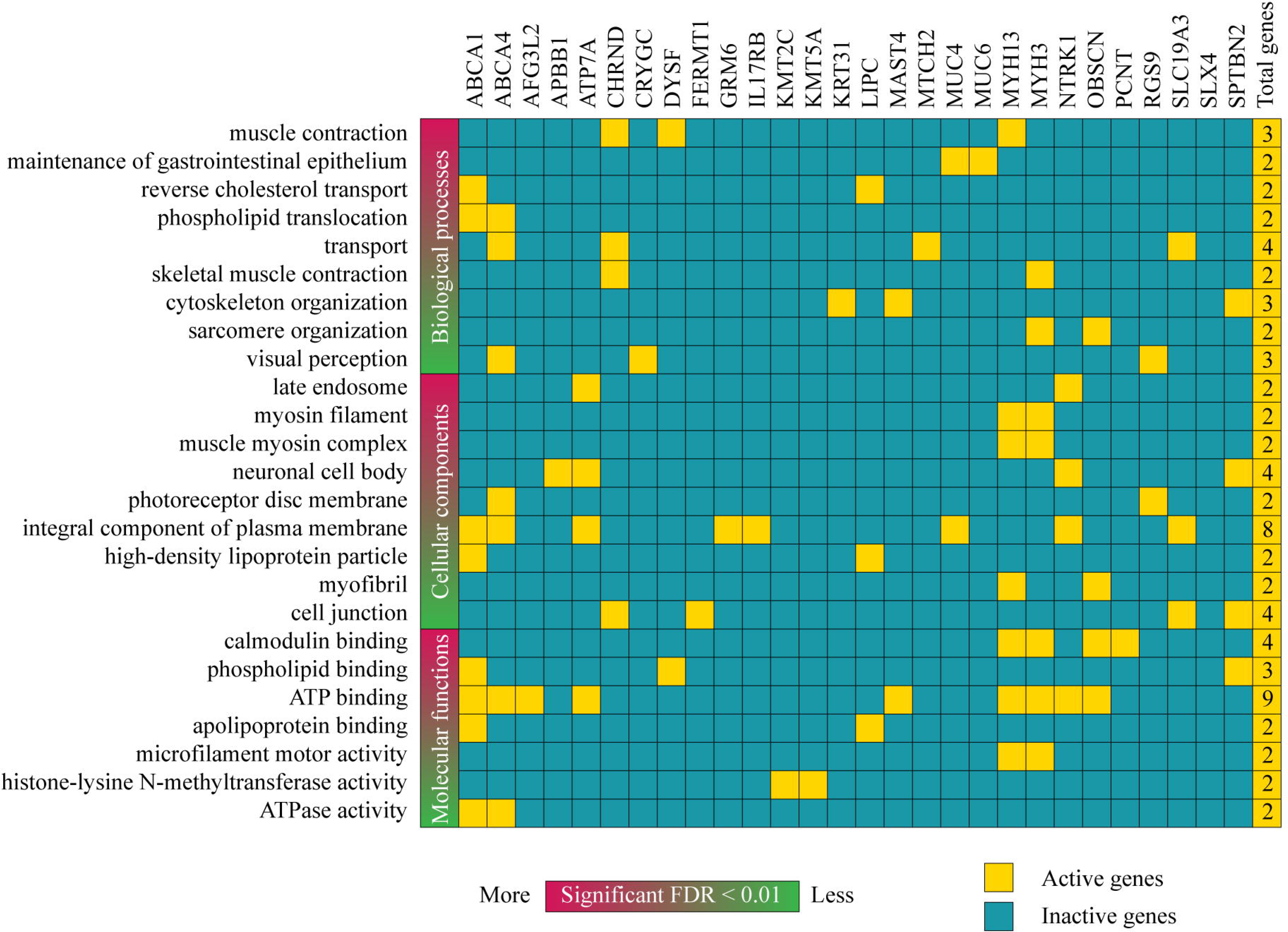
NTRKI mutations. a) Pattern of autosomal recessive inheritance of CIPA patient with respect to the pathogenic mutation rs80356677 (Arg674Tyr). b) NTRK1 protein structure showing the location of Arg 602 and Arg 674 mutations.

### Genomic DNA analysis and the pain matrix genes

After performing the genomic DNA analysis using the TruSight One (TSO) Next-Generation Sequencing (NGS) Panel (Illumina, Inc. San Diego, CA, USA), it is important to mention that the total aligned reads was 14,356,459 (father), 17,975032 (mother) and 20,225,887 (patient). The percentage of reads passing filter that aligned to the reference was 99.9% for all samples. The percentage of targets with coverage greater than 20X was 27.3% for the father, 30.3% for the mother and 30.4% for the patient with CIPA. Additionally, the analysis of 4,811 genes and 18,933 variants was performed in the BaseSpace Variant Interpreter software (Illumina).

In the first step, 12,068 variants (pathogenic, likely pathogenic, VUS, likely benign and benign variants) passed the small variant QC metrics, copy number QC metrics and structural variant QC metrics. In the second step, only pathogenic, probably pathogenic and VUS were taken into account according to the BaseSpace Variant Interpreter, obtaining 384 variants of 233 genes. In the third step, only deleterious, damaging, probably damaging and VUS were taken into account according to the SIFT and PolyPhen-2 bioinformatics tools(38,39), obtaining 178 variants of 133 genes. In the fourth step, a manual curation of each variant was performed to confirm its clinical significance through the ClinVar and LOVD(35,40), remaining in total 68 pathogenic / likely pathogenic variants (Table 2), and 76 VUS in 105 genes. Of these 144 variants, 71 were missense variants, 2 were missense variants / splice region variants, 3 were frameshift variants, 13 were inframe deletions, 12 were inframe insertions, 7 were intron variants / non-coding transcript variants, 1 was non-coding transcript exon variant, 3 were splice donor variants, 19 were splice region variants / intron variants, 1 was splice region variant / non-coding transcript exon variant, 3 were splice region variants / synonymous variants, 8 were stop gained variants, and 1 was stop gained / splice region variant (S2 Table).

**Table 2.**
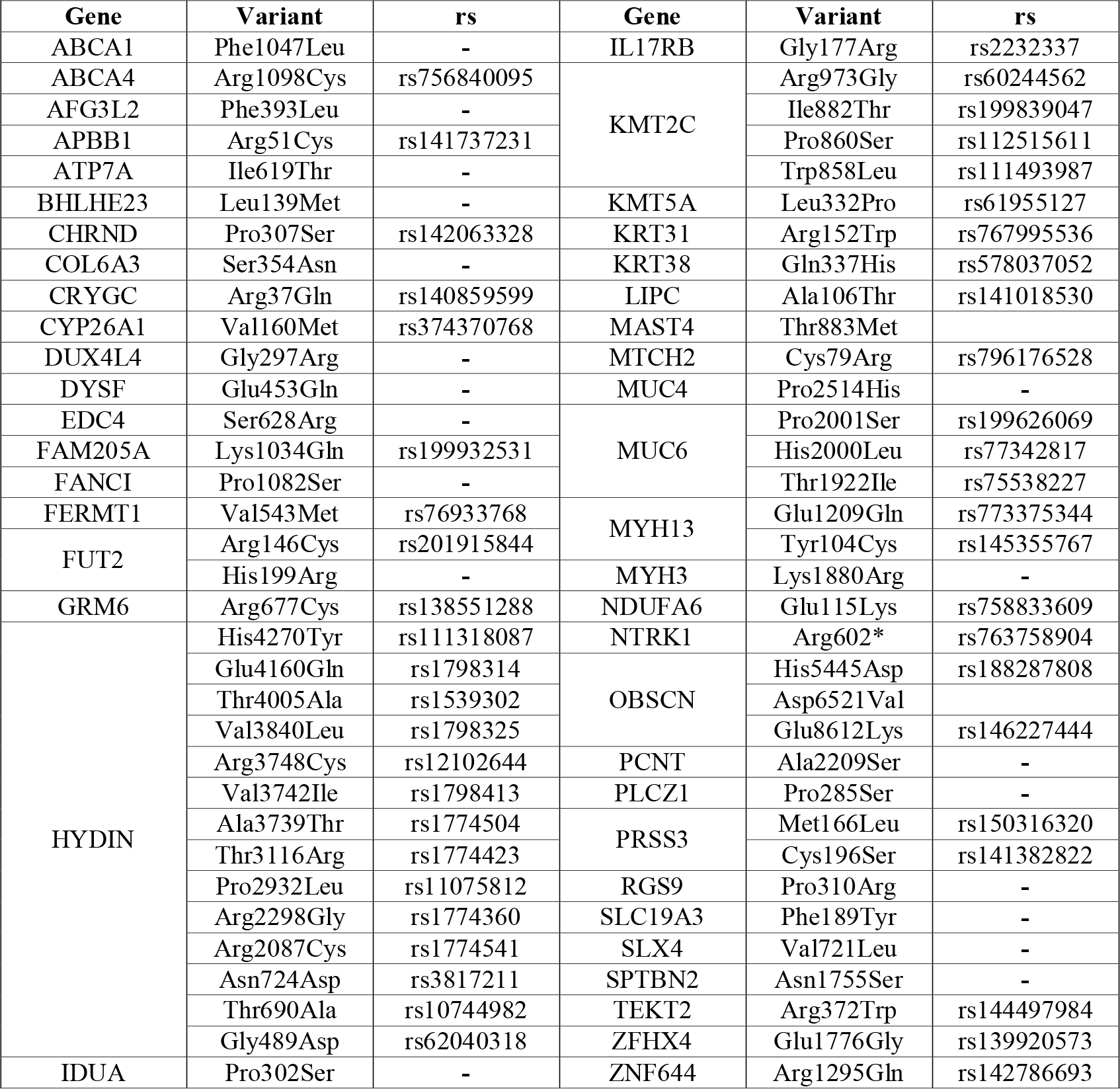
Pathogenic and likely pathogenic variants in the CIPA patient after DNA genomic analysis.

The enrichment analysis of GO terms related to biological processes, cellular components and molecular functions was carried on in 45 genes with pathogenic and likely pathogenic variants using DAVID Bioinformatics Resource(41). Only 28 of 45 genes were involved in almost one of the categories showed as a heatmap in Fig 4. The most significant biological processes (BP) with a false discovery rate (FDR) < 0.01 were muscle contraction, maintenance of gastrointestinal epithelium, reverse cholesterol transport, phospholipid translocation, skeletal muscle contraction, cytoskeleton organization, sarcomere organization and visual perception. The BPs with the highest number of active genes were transport, muscle contraction and cytoskeleton organization. The most significant cellular components (CC) with a FDR < 0.01 were late endosome, myosin filament, muscle myosin complex, neuronal cell body, photoreceptor disc membrane, integral component o plasma membrane, high-density lipoprotein particle, myofibril and cell junction. The CCs with the highest number of active genes were integral component of plasma membrane and neural cell body. In addition, the most significant molecular functions (MF) with a FDR < 0.01 were calmodium binding, phospholipid binding, ATP binding, apolipoprotein binding, microfilament motor activity, histone-lysine N-methyltransferase activity and ATPase activity(41). The MFs with the highest number of active genes were ATP binding and calmodium binding (Fig 4). The function of all these genes is detailed in the S3 Table.

**Fig 4.**
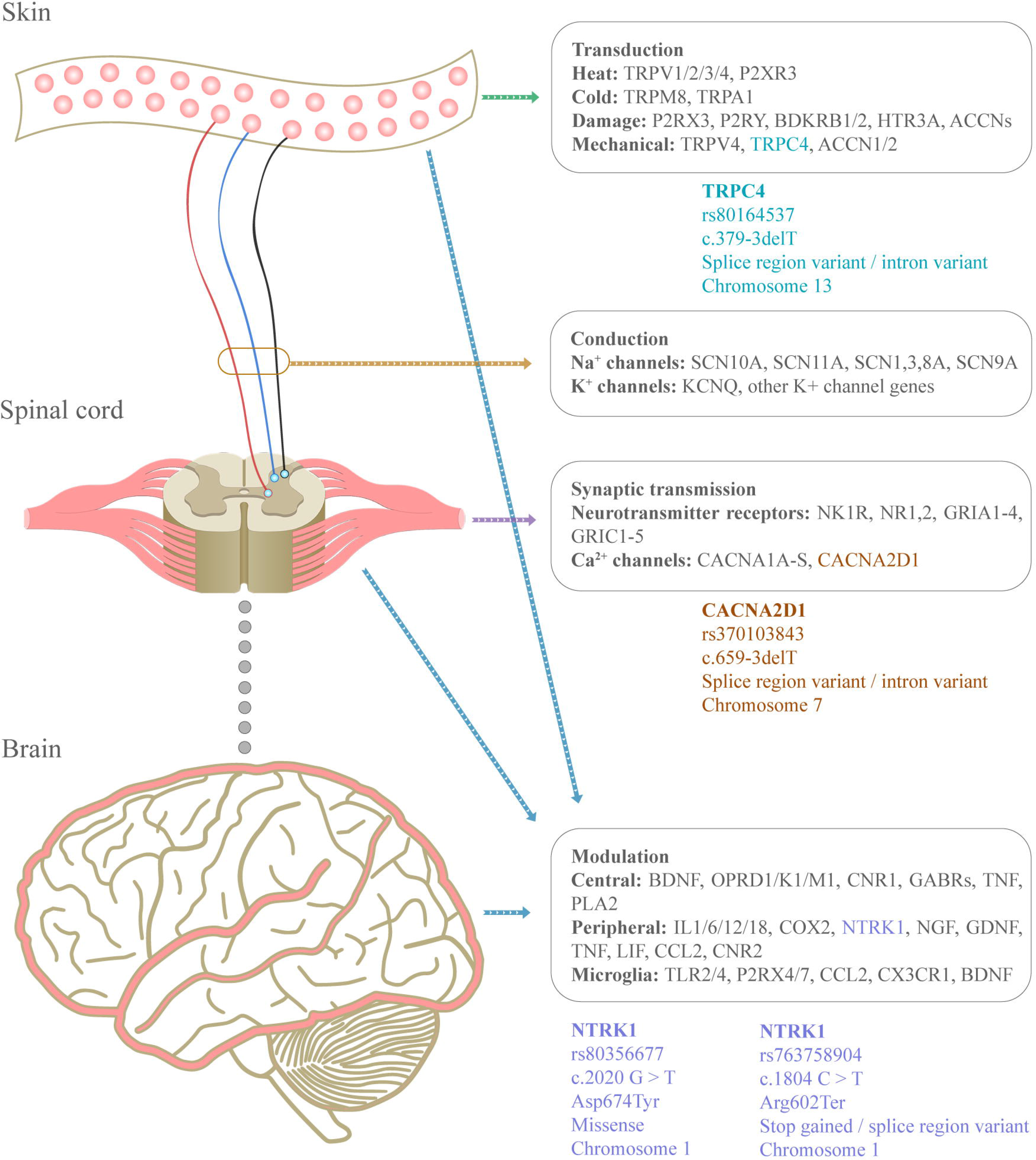
Heatmap of active and inactive genes with pathogenic and/or likely pathogenic variants involved in biological processes, cellular components and molecular functions according to the GO enrichment analysis.

After the DNA genomic analysis performed on CIPA patient, three pain matrix genes presented four variants. NTRK1, which acts on the modulation of the nervous stimulus, presented two pathogenic variants: 1) rs80356677 (c.2020 G>T) is a pathogenic missense variant located on chromosome 1 and consists of the change of Asp to Tyr at amino acid 674; and 2) rs763758904 (c.1804 C>T) is a pathogenic stop gained / splice region variant located on chromosome 1 and consists of the change of Arg to a stop codon. In regard to the protein structure of NTRK1, the rs763758904 (Arg602*) mutation is located in the α-D-helix (Fig 3b). The CACNA2D1 gene, which acts in the synaptic transmission through Ca^2+^ channels, presented the VUS rs370103843 (c.659-3delT) that is a splice region variant / intron variant located on chromosome 7 and consists of a TTT deletion. Lastly, the TRPC4 gene, which acts on the mechanical transduction of the nervous stimuli, presented the VUS rs80164537 (c.379-3delT) that is a splice region variant / intron variant located on chromosome 13 and consists of a TTT deletion (Fig 5). The full list of the pain matrix genes is detailed in the S4 Table.

**Fig 5.**
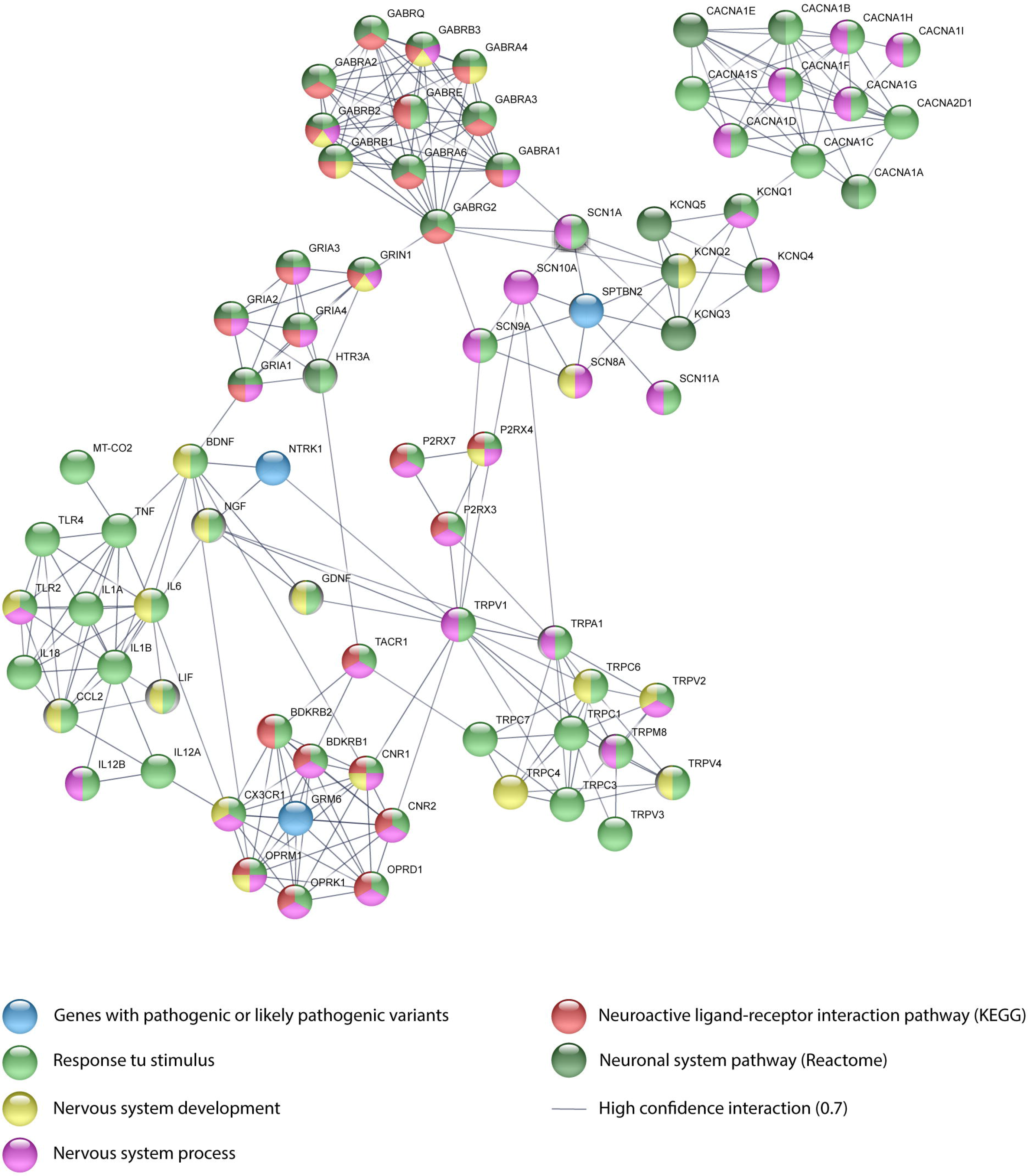
The pain matrix genes.

### Networking and the pain matrix genes

Fig 6 shows a PPi network with a high confidence of 0.7 created using String Database(42). This network was made up of known and predicted interactions between genes with pathogenic and/or likely pathogenic variants and the pain matrix genes. As results, three genes with pathogenic and/or likely pathogenic variants (NTRK1, SPTBN2 and GRM6) were associated with several genes of the pain matrix. NTRK1 interacts with NGF, BDNF and TRPV1. GRM6 interacts with CNR1, DPRD1, CNR2, BDKRB2, OPRK1, BDKRB1, OPRM1 and CX3CR1. Lastly, SPTBN2 interacts with SCN1A, KCNQ3, KCNQ2, SCN11A, SCN10A, SCN8A and SCN9A.

**Fig 6.**
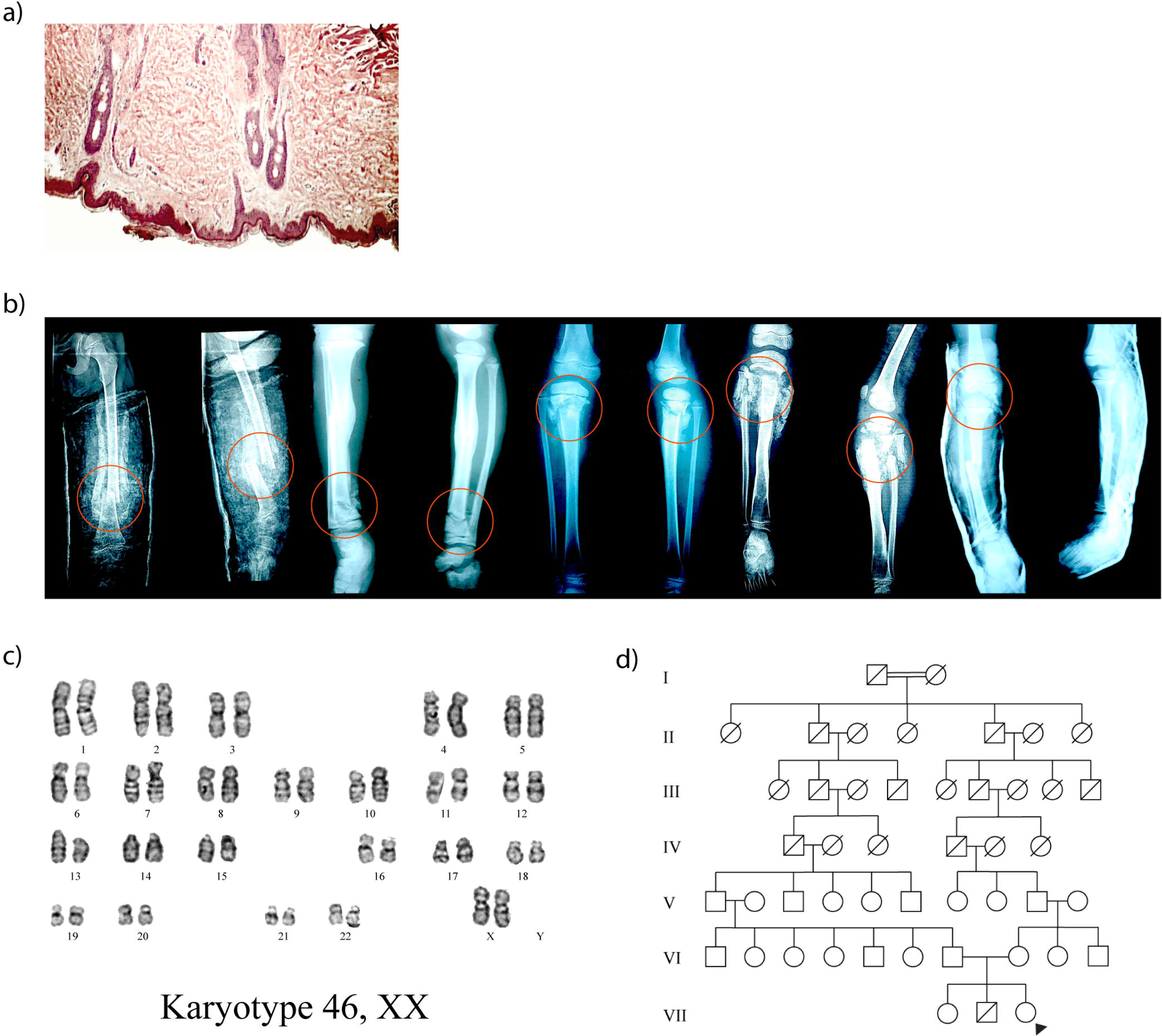
Protein-protein interaction network between the pain matrix genes and genes with pathogenic or likely pathogenic variants.

## DISCUSSION

CIPA (HSAN-IV) is a painless severe genetic disorder associated with overwhelming complications, often leading to lethal consequences(1). Since the first report on this pathology mentioned by Dearborn in 1932(13) and published by Swanson in 1963(14), only a few hundred of cases have been reported worldwide. Each of these publications has provided a significant contribution to better understand the clinical and genetic aspects of this autosomal recessive disorder(8,12,23–28,30,31,15–22).

In our study, we investigated one Native American patient from a family who lives in a high-altitude indigenous community (2,418 MASL) located north of the Ecuadorian highlands. Regarding the analysis of ancestry informative markers, the Native American ancestry of patient was the most prevalent with 87.8% followed by African ancestry with 6.6% and European ancestry with 5.6% (Fig 2a-c). Therefore, this is the first case from the high-altitude Ecuadorian indigenous population reported worldwide.

Nine years of clinical record have shown that the patient with CIPA has presented several health problems such as bone fractures, self-mutilation, osteochondroma, intellectual disability, Riga-Fede disease, ulcers and fever. In addition, it is important to mention that the consanguinity of their ancestors is an important factor for the patient to have this autosomal recessive disease (Fig 1a-e).

The NTRK1 pathway is responsible for innervating skin through sensory axons, as well being essential for the survival of pain receptors and maintenance of autonomic sympathetic postganglionic neurons(43). Through this pathway, sympathetic neurons regulate sweating, piloerection and vasoconstriction in hot and cold environments. NTRK1 pathway also involves endocytosis and vesicular transport carried through endosomes that are being transported to the cell body for activating the signaling molecule that promotes neural differentiation(43). NTRK1, a transmembrane glycosylated protein of 140 kDa is composed of the extracellular domain, the intracellular domain, the tyrosine kinase domain and a carboxyl terminal tail. The encoded NTRK1 protein, a member of the tyrosine kinase family, binds to a NGF receptor stimulating homodimer formation and autophosphorylation of tyrosine residues(44), serving as anchors for binding to downstream signaling molecules. However, with altered NTRK1, the NFG protein leads to aberrations in processing and survival of sympathetic ganglion neurons and pain receptors(45). After carrying out the mutational analysis of NTRK1 and the genomic DNA analysis, the Ecuadorian indigenous patient presented two pathogenic mutations in the kinase domain of NTRK1. rs80356677 (c.2020 G>T; Asp674Tyr) is a pathogenic missense variant characterized by the change of Asp to Tyr in the amino acid 674. Parents presented the heterozygous genotype (G/T), while the patient presented the homozygous mutant genotype (T/T). This genetic variant has been reported by Indo *et al* (2001)(20). According to the 1000 Genomes Project (Phase 3) and/or the Exome Aggregation Consortium (ExAC), rs80356677 has a minor allele (T) frequency < 0.005 in the East Asian population (Japan) and 0.000 in other populations worldwide(46,47). On the other hand, rs763758904 (c.1804 C>T; Arg602*) is a pathogenic stop gained / splice region variant characterized by the change of Arg to stop codon. Parents presented the heterozygous genotype (C/T), while the patient presented the homozygous mutant genotype (T/T). This genetic variant has been reported by Wang *et al* (2016) in a study on five Chinese children with CIPA. According to the UK10K Project, rs763758904 has a minor allele (T) frequency < 0.0003 in the United Kingdom and 0.0000 in other populations worlwide(48).

The pain matrix shown in Fig 5 is a complex circuit of pain perception proposed by Foulkes *et al* (2008)(11). Different genes/proteins are involved in the four phases of the pain matrix that are transduction, conduction, synaptic transmission and modulation. In addition to the pathogenic variants of NTRK1, the patient with CIPA presented two VUS in CACNA2D1 (rs370103843) and TRPC4 (rs80164537) genes. CACNA2D1 acts in the synaptic transmission through Ca^2+^ channels. rs370103843 (c.659-3delT) is a splice region variant / intron variant located on chromosome 7 and consists of a TTT deletion. TRPC4 acts on the mechanical transduction of the nervous stimuli. rs80164537 (c.379-3delT) is a splice region variant / intron variant located on chromosome 13 and consists of a TTT deletion. The presence of both variants could be associated with the alteration of the patient’s perception of pain. According to the ExAC, the global minor allele frequency of rs370103843 is 0.075, and of rs80164537 is 0.383(47).

On the other hand, the DNA genomic analysis through next-generation sequencing showed that the CIPA patient presented 68 pathogenic and/or likely pathogenic variants in 45 genes (Table 2). After an enrichment analysis of GO(41), only 28 genes were involved in almost one of the categories showed as a heatmap in Fig 4. The most significant BP was muscle contraction, and transport was the BP with more active genes (ABCA4, CHRND, MTCH2 and SLC19A3) with pathogenic and/or likely pathogenic variants. The most significant CC was late endosome, and integral component of plasma membrane was the CC with more active genes (ABCA1, ABCA4, ATP7A, GRM6, IL17RB, MUC4, NTRK1 and SLC19A3) with pathogenic and/or likely pathogenic variants. The most significant MF was calmodium binding, and ATP binding was the MF with more active genes (ABCA1, ABCA4, AFG3L2, ATP7A, MAST4, MYH13, MYH3, NTRK1 and OBSCN) with pathogenic and/or likely pathogenic variants. Lastly, the GO terms where NTRK1 was active were late endosome, neural cell body, integral component of plasma membrane and ATP binding. The bioinformatics analysis of the Ecuadorian patient serves as a guide to better understand the possible symptoms and complications of CIPA patients worldwide.

In regard to the networking analysis, the PPi between genes with pathogenic and/or likely pathogenic variants and the pain matrix genes demonstrates that NTRK1 interacts with NGF, BDNF and TRPV1. NGF and BDNF are involved in nervous system development and response to stimulus. TRPV1 is involved in nervous system process and response to stimulus. GRM6 interacts with CNR1, DPRD1, CNR2, BDKRB2, OPRK1, BDKRB1, OPRM1 and CX3CR1. OPRK1, OPRD1, CNR2 and BDKRB1 are involved in nervous system process, response to stimulus and neuroactive ligand-receptor interaction pathway. OPRM1 and CNR1 are involved in nervous system process, response to stimulus, neuroactive ligand-receptor interaction pathway and nervous system development. BDKRB2 is involved in neuroactive ligand-receptor interaction pathway and response to stimulus. CX3CR1 is involved in nervous system process, response to stimulus and nervous system development. Lastly, SPTBN2 interacts with SCN1A, KCNQ3, KCNQ2, SCN11A, SCN10A, SCN8A and SCN9A. SCN11A, SCN1A and SCN9A are involved in nervous system process and response to stimulus. SCN8A is involved in nervous system development and process. SCN10A is involved in nervous system process. KCNQ2 is involved in nervous system development and the neuronal system pathway. Lastly, KCNQ3 is involved in the neural system pathway(41,49,50).

This PPi network can contribute to understand how different genes with pathogenic variants influence the development of symptoms and health problems in patients with CIPA worldwide.

In conclusion, we conducted for the first time clinical, ancestral, genomics, PPi networking and GO enrichment analyses in a high-altitude Native American (indigenous) Ecuadorian patient with CIPA and with family history of consanguinity, whose results were associated with the pain matrix gene in order to find new genes associated with the pathogenesis of this complex and infrequent genetic disease.

## MATERIALS AND METHODS

### Biological samples

Skin biopsy of sternal region was taken to analyze presence or absence of nerve terminals by immunohistochemical staining for S100 protein. Additionally, peripheral blood samples were extracted from CIPA patient and her parents using FTA buffer (GE Healthcare, UK) in order to perform genomic, ancestral and cytogenetic analyses. The present study was approved by the Bioethics Committee from Universidad de las Américas (No. 2015-0702), and all individuals signed their respective informed consent.

### DNA extraction and purification

Genomic DNA was extracted from whole blood using the PureLink Genomic DNA Kit (Invitrogen, Carlsbad, CA), and purified using Amicon Ultra centrifugal filters (Merck, Darmstadt, Germany). Genomic DNA of individuals presented a concentration of 45 ng/μL (mother), 27 ng/μL (father) and 36 ng/μL (patient). This calculation was obtained using a Qubit 4 (Thermo Scientific, Waltham, MA).

### Ancestry informative markers

Ancestry informative insertion/deletion markers (AIM-INDELs) have been used as a tool to fully understand ancestry and geographical differentiation of CIPA patient’s family. We selected 3 samples self-identified as indigenous from a high-altitude community located in the Ecuadorian highlands region at 2,418 MASL.

The amplification was done in one multiplex reaction with the same 46 AIM-INDELs primers and procedure according to Pereira *et al* (2012) and Zambrano *et al* (2017)(51,52). Fragment separation and detection were performed in ABI and 3500 genetic analyzers (Applied Biosystems, Austin, TX), collected with Data Collection v3.1 and analyzed by Gene Mapper Software 5 (Applied Biosystems).

Data was analyzed using Structure v.2.3.4 software for the ancestral proportions using Africans, Europeans and Native Americans as reference populations (population selected because historical records); and lastly, RStudio v.1.1.453 for the principal component analysis (PCA)(53).

### Karyotype

800 μl of peripheral blood were cultured in 5 ml of RPMI 1640 medium supplemented with fetal calf serum, L-glutamine, antibiotic-antimycotic, hepes-buffer and phytohemagglutinin (Gibco, Grand Island, NY). After 72 hours of incubation at 37 ºC, 200 μl of Colcemid (Gibco) was added and kept in for further 40 min. Cells were exposed to hypotonic shock with potassium chloride 0.075 M for 25 min at 37 °C and fixed with Carnoy’s solution at least three times (3:1 v/v methanol and glacial acetic acid)(54). The pellet was resuspended in 1 ml of Carnoy’s solution and spread in coded glass slides. After overnight incubation at 56 ºC, chromosomes were G-banded using trypsin and stained with Giemsa. Slides were analyzed at 1000X magnification in a conventional microscope (Nikon Eclipse E600, Nikon, Japan). By using GenASIs BandView^®^, metaphases with chromosomes were scored.

### Mutational analysis of NTRK1

NTRK1 genotyping was performed using DNA sequencing analysis through the Sanger method. A final volume of 50 μL was used for each PCR reaction for the 54 SNPs located into 14 regions. Each reaction consisted of 34 μL of Milli-Q water, 4 μL of DNA template (20 ng/μL), 0.2 μM of each deoxynucleotide triphosphate (dNTPs), 1.5 mM of MgCl_2_, 2.5 U of Taq DNA polymerase, 5 μL of 10× buffer (500 mM of KCl, 200 mM of Tris-HCl, pH=8.4), and 0.4 μM of forward (FW) and reverse (RV) primers. Table 1 details primer sequence, weight fragment, exon location, reference sequence, allele change, residual change, clinical significance, function of all 54 variants analyzed. The PCR program for all genetic variants started with an initial denaturation stage lasting 5 min at 95 °C, followed by 35 cycles of 30 s at 94 °C, 45 s at 57.4 °C, 45 s at 72 °C, and a final elongation for 3 min at 72 °C. Each run was completed using a Sure Cycler 8800 thermocycler (Agilent, Santa Clara, CA). The amplified fragments were analyzed by electrophoresis in 2% agarose gels with SYBR Safe DNA gel stain (Invitrogen, Carlsbad, CA), and was observed in an Enduro™ GDS Touch transilluminator (Labnet International Inc, Edison, NJ).

**Table 1.**
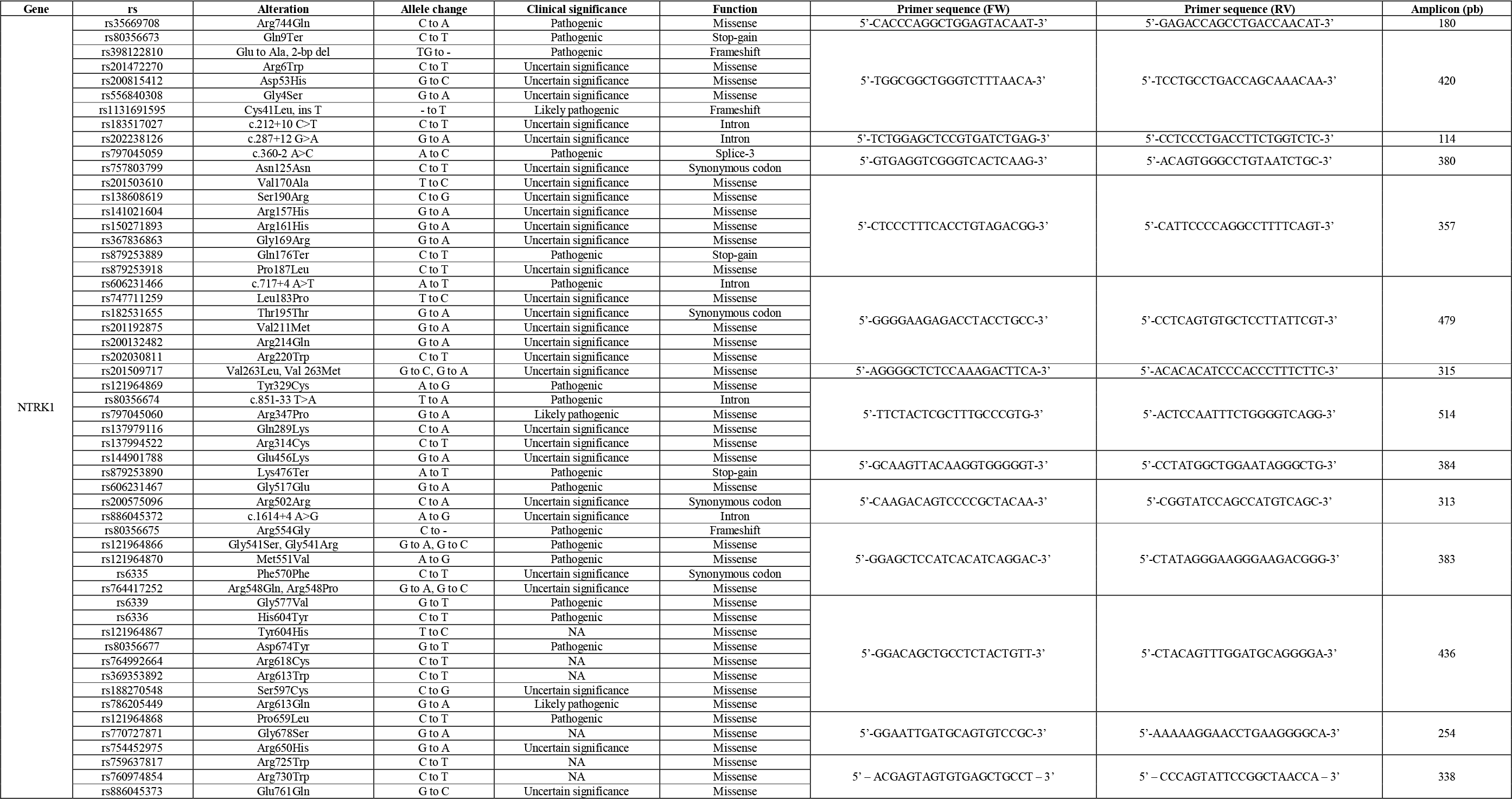
Studied mutations of the NTRK1 gene, and their PCR conditions.

Subsequently, the PCR amplicons were analyzed through sequence analysis, using the Genetic Analyzer 3500 (Applied Biosystems, Austin, TX). The final volume of the reaction was 19.6 μL and contained 5.6 μL of Milli-Q water, 4 μL of 5X buffer, 2 μL of primers FW or RV (3.2 pmol), 2 μL of BigDye Terminator v3.1 sequencing standard (Applied Biosystems, Austin, TX), and 6 μL of PCR product (3 to 10 ng). Once the product was amplified, it was purified using QIAquick PCR Purification Kit (Qiagen, Chatsworth, CA). The amplicon program consisted of 3 min at 96 °C, followed by 30 cycles of 10 s at 96 °C, 5 s at 50 °C, and 4 min at 60 °C. Finally, sequence analysis was performed using 3500 Data Collection Software v3.1 (Applied Biosystems) and the alignment with sequence from GeneBank (NTRK1 NG_007493.1) was performed using Sequencing Analysis Software v6.0 (Applied Biosystems) and CLC Main Workbench 8.0 (CLC bio, Aarhus, Denmark).

### Genomic DNA analysis

Genomic DNA of three samples (mother, father and patient) was enriched by using the TruSight One (TSO) Next-Generation Sequencing (NGS) Panel (Illumina, Inc. San Diego, CA, USA), which includes 125,395 probes targeting a 12-Mb region spanning ~62,000 target exons of 4,811 genes, and sequenced on the Illumina MiSeq platform. Raw sequence reads were processed and aligned against the human NCBI GRCh37/hg19 reference genome assembly using the BWA software. The 80-mer probes target libraries with ~500 bp mean fragment sizes and ~300 bp insert sizes, enriching a broad footprint of 350-650 based centered symmetrically around the midpoint of the probe. Therefore, in addition to covering the main exon regions, the panels cover exon-flanking regions, which can provide important biological information such as splice sites or regulatory regions. The TSO coverage is ≥20× on 95% of the target regions in the panel. The TSO full gene list is detailed in the S1 Table, and the sequencing panel workflow that consists of tagment genomic DNA, clean up tagmented DNA, amplify tagmented DNA, clean up amplified DNA, hybridize probes, capture hybridized probes, perform second hybridization, perform second capture, clean up captured library, amplify enriched library, clean up amplified enriched library and check enriched library can be found in the manufacturer’s website (https://support.illumina.com/content/dam/illumina-support/documents/documentation/chemistry_documentation/trusight_one/trusight-one-sequencing-panel-reference-guide-15046431-03.pdf).

To analyze and characterize data generated from targeted sequencing, the following software and databases were implemented in our bioinformatics pipeline: the analysis of alignment and annotation was done with the BWA software. The BaseSpace Variant Interpreter software (Illumina) imports and annotates variant call files in the VCF file format generated during analysis of sequencing data. After import, BaseSpace Interpreter provides commands to filter results using various filtering options, and classifying variants according to their clinical significance (pathogenic, likely pathogenic, variants of uncertain significance (VUS), likely benign and benign variants) (https://support.illumina.com/sequencing/sequencing_software/basespace-variant-interpreter.html). SIFT predicts whether an amino acid substitution affects protein function based on sequence homology and the physical properties of amino acids (http://sift.bii.a-star.edu.sg/)(39). PolyPhen-2 is a tool that predicts possible impact of an amino acid substitution on the structure and function of a human protein using straightforward physical and comparative considerations (http://genetics.bwh.harvard.edu/pph2/)(38). ClinVar is a public archive of reports of the relationships among human variations and phenotypes (https://www.ncbi.nlm.nih.gov/clinvar/)(35). The Human Gene Mutation Database constitutes a comprehensive collection of published germline mutations in nuclear genes that underlie, or are closely associated with human inherited disease (http://www.hgmd.cf.ac.uk/ac/index.php)(36). Leiden Open Variation Databases (LOVD) provides a flexible tool for gene-centered collection and display of DNA variations (http://www.lovd.nl/3.0/home)(40). The database for annotation, visualization and integrated discovery (DAVID) provides a comprehensive set of functional annotation tool to understand biological meaning behind large list of genes. Lastly, the enrichment analysis of GO terms related to biological processes, cellular components and molecular functions was done with DAVID (https://david.ncifcrf.gov/)(41).

### Protein-protein interaction (PPi) networking and the pain matrix genes

A correlation of the filtered pathogenic, likely pathogenic and VUS variants was performed with the pain matrix genes proposed by Foulkes *et al* (2008)(11), in order to better understand CIPA pathogenesis. In addition, a protein-protein interaction network with a high confidence of 0.7 was created using String Database(42). This network was made up of known and predicted interactions between genes with pathogenic and/or likely pathogenic variants and the pain matrix genes.

## Acknowledgements

Universidad UTE supported this research.

## Author contributions

ALC conceived the subject and wrote the paper. AKZ and PGR performed the next-generation sequencing. AKZ performed the ancestral analysis. BAE, EC and GPM made substantial contribution with the clinical data of the patient. SG performed the NTRK1 protein structure analysis. APV performed the cytogenetics analysis. IAC, JGC, VY, PEL did data curation and supplementary data. CPyM supervised the research project. Lastly, all authors reviewed and/or edited the article before submission.

## Data availability statement

All data generated or analyzed during this study are included in this published article and its Supplementary Dataset.

## REFERENCES

1. Altassan R, Saud H Al, Masoodi TA, Dosssari H Al, Khalifa O, Al-Zaidan H, et al. Exome sequencing identifies novel NTRK1 mutations in patients with HSAN-IV phenotype. Am J Med Genet Part A. 2017;

2. Axelrod FB, Gold-Von Simson G. Hereditary sensory and autonomic neuropathies: Types II, III, and IV. Orphanet J Rare Dis. 2007;

3. Indo Y. Nerve growth factor and the physiology of??pain: Lessons from congenital insensitivity to pain with anhidrosis. Clinical Genetics. 2012.

4. Kilic SS, Ozturk R, Sarisozen B, Rotthier A, Baets J, Timmerman V. Humoral immunodeficiency in congenital insensitivity to pain with anhidrosis. Neurogenetics. 2009;

5. Bonkowsky JL, Johnson J, Carey JC, Smith a G, Swoboda KJ. An infant with primary tooth loss and palmar hyperkeratosis: a novel mutation in the NTRK1 gene causing congenital insensitivity to pain with anhidrosis. Pediatrics. 2003;112(3):e237–41.

6. Sayyahfar S, Chavoshzadeh Z, Khaledi M, Madadi F, Yeganeh MH, Sawamura D, et al. Congenital insensitivity to pain with anhidrosis presenting with palmoplantar keratoderma. Pediatr Dermatol. 2013;

7. Barone R, Lempereur L, Anastasi M, Parano E, Pavone P. Congenital insensitivity to pain with anhidrosis (NTRK1 mutation) and early onset renal disease: Clinical report on three sibs with a 25-year follow-up in one of them. Neuropediatrics. 2005;

8. Pérez-López LM, Cabrera-González M, Gutiérrez-de la Iglesia D, Ricart S, Knörr-Giménez G. Update review and clinical presentation in congenital insensitivity to pain and anhidrosis. Case Rep Pediatr. 2015;2015:1–7.

9. Daneshjou K, Jafarieh H, Raaeskarami S-R. Congenital Insensitivity to Pain and Anhydrosis (CIPA) Syndrome; A Report of 4 Cases. Iran J Pediatr. 2012;22(3):412–6.

10. Tracey I, Mantyh PW. The Cerebral Signature for Pain Perception and Its Modulation. Neuron. 2007.

11. Foulkes T, Wood JN. Pain Genes. PLoS Genet. 2008;4(7):e1000086.

12. Gao L, Guo H, Ye N, Bai Y, Liu X, Yu P, et al. Oral and Craniofacial Manifestations and Two Novel Missense Mutations of the NTRK1 Gene Identified in the Patient with Congenital Insensitivity to Pain with Anhidrosis. PLoS One. 2013;8(6).

13. Van Ness Dearborn G. A case of congenital general pure analgesia. J Nerv Ment Dis. 1932;

14. Swanson AG. Congenital Insensitivity to Pain with Anhydrosis: A Unique Syndrome in Two Male Siblings. Arch Neurol. 1963;

15. Tuncbilek G, Oztekin C, Kayikcioglu A. Calcaneal ulcer in a child with congenital insensitivity to pain syndrome. Scand J Plast Reconstr Surg Hand Surg. 2005;

16. Indo Y, Tsuruta M, Hayashida Y, Karim M, Ohta K, Kawano T, et al. Mutations in the TRKA/NGF receptor gene in patients with congenital insensitivity to pain with anhidrosis. Nat Genet. 1996;13(4):485–8.

17. Grills BL, Schuijers JA. Immunohistochemical localization of nerve growth factor in fractured and unfractured rat bone. Acta Orthop Scand. 1998;

18. Indo Y. Molecular basis of congenital insensitivity to pain with anhidrosis (CIPA): Mutations and polymorphisms in TRKA (NTRK1) gene encoding the receptor tyrosine kinase for nerve growth factor. Vol. 18, Human Mutation. 2001. p. 462–71.

19. Jarade EF, El-Sheikh HF, Tabbara KF. Indolent corneal ulcers in a patient with congenital insensitivity to pain with anhidrosis: A case report and literature review. Eur J Ophthalmol. 2002;

20. Indo Y, Mardy S, Miura Y, Moosa A, Ismail EAR, Toscano E, et al. Congenital insensitivity to pain with anhidrosis (CIPA): Novel mutations of the TRKA (NTRK1) gene, a putative uniparental disomy, and a linkage of the mutant TRKA and PKLR genes in a family with CIPA and pyruvate kinase deficiency. Hum Mutat. 2001;18(4):308–18.

21. Weier HUG, Rhein AP, Shadravan F, Collins C, Polikoff D. Rapid physical mapping of the human trk protooncogene (NTRK1) to human chromosome 1q21-q22 by P1 clone selection, fluorescence in situ hybridization (FISH), and computer-assisted microscopy. Genomics. 1995;26(2):390–3.

22. Miura Y, Mardy S, Awaya Y, Nihei K, Endo F, Matsuda I, et al. Mutation and polymorphism analysis of the TRKA (NTRK1) gene encoding a high-affinity receptor for nerve growth factor in congenital insensitivity to pain with anhidrosis (CIPA) families. Hum Genet. 2000;106(1):116–24.

23. Lin YP, Su YN, Weng WC, Lee WT. Novel neurotrophic tyrosine kinase receptor type 1 gene mutation associated with congenital insensitivity to pain with anhidrosis. J Child Neurol. 2010;

24. Mardy S, Miura Y, Endo F, Matsuda I, Indo Y. Congenital insensitivity to pain with anhidrosis (CIPA): effect of TRKA (NTRK1) missense mutations on autophosphorylation of the receptor tyrosine kinase for nerve growth factor. Hum Mol Genet. 2001;

25. Schreiber R, Levy J, Loewenthal N, Pinsk V, Hershkovitz E. Decreased first phase insulin response in children with congenital insensitivity to pain with anhidrosis. J Pediatr Endocrinol Metab. 2005;

26. Brandes IF, Stuth EAE. Use of BIS monitor in a child with congenital insensitivity to pain with anhidrosis. Paediatr Anaesth. 2006;

27. Indo Y. Genetics of congenital insensitivity to pain with anhidrosis (CIPA) or hereditary sensory and autonomic neuropathy type IV. Clin Auton Res. 2002;

28. Tanaka M, Sotomatsu A, Kanai H, Hirai S. [Iron-dependent cytotoxic effects of dopa on cultured neurons of the dorsal root ganglia]. Rinsho Shinkeigaku. 1990;

29. Melamed I, Levy J, Parvari R, Gelfand EW. A novel lymphocyte singnaling defect: trk a mutation in the syndrome of congenital insensitivity to pain and anhidrosis (CIPA). J Clin Immunol. 2004;24(4):441–8.

30. Abdulla M, Khaled SS, Khaled YS, Kapoor H. Congenital insensitivity to pain in a child attending a paediatric fracture clinic. J Pediatr Orthop Part B. 2014;

31. Franco ML, Melero C, Sarasola E, Acebo P, Luque A, Calatayud-Baselga I, et al. Mutations in TrkA causing congenital insensitivity to pain with anhidrosis (CIPA) induce misfolding, aggregation, and mutation-dependent neurodegeneration by dysfunction of the autophagic flux. J Biol Chem. 2016;

32. Varma AV, McBride L, Marble M, Tilton A. Congenital insensitivity to pain and anhidrosis: Case report and review of findings along neuro-immune axis in the disorder. J Neurol Sci. 2016;

33. Zhang Y-Z, Moheban DB, Conway BR, Bhattacharyya A, Segal RA. Cell surface Trk receptors mediate NGF-induced survival while internalized receptors regulate NGF-induced differentiation. J Neurosci. 2000;

34. Indo Y. Neurobiology of pain, interoception and emotional response: Lessons from nerve growth factor-dependent neurons. Eur J Neurosci. 2014;

35. Landrum MJ, Lee JM, Riley GR, Jang W, Rubinstein WS, Church DM, et al. ClinVar: Public archive of relationships among sequence variation and human phenotype. Nucleic Acids Res. 2014;42(D1).

36. Stenson PD, Mort M, Ball E V., Evans K, Hayden M, Heywood S, et al. The Human Gene Mutation Database: towards a comprehensive repository of inherited mutation data for medical research, genetic diagnosis and next-generation sequencing studies. Human Genetics. 2017.

37. Indo Y, Tsuruta M, Hayashida Y, Karim MA, Ohta K, Kawano T, et al. Mutations in the TRKA/NGF receptor gene in patients with congenital insensitivity to pain with anhidrosis. Nat Genet. 1996;13(4):485–8.

38. Adzhubei I, Jordan DM, Sunyaev SR. Predicting functional effect of human missense mutations using PolyPhen-2. Curr Protoc Hum Genet. 2013;

39. Ng PC, Henikoff S. SIFT: Predicting amino acid changes that affect protein function. Nucleic Acids Res. 2003;

40. Fokkema IFAC, Taschner PEM, Schaafsma GCP, Celli J, Laros JFJ, den Dunnen JT. LOVD v.2.0: The next generation in gene variant databases. Hum Mutat. 2011;

41. Huang DW, Sherman BT, Lempicki RA. Systematic and integrative analysis of large gene lists using DAVID bioinformatics resources. Nat Protoc. 2009 Dec;4(1):44–57.

42. Szklarczyk D, Franceschini A, Wyder S, Forslund K, Heller D, Huerta-Cepas J, et al. STRING v10: protein-protein interaction networks, integrated over the tree of life. Nucleic Acids Res. 2015 Jan 28;43(Database issue):D447–52.

43. Beigelman A, Levy J, Hadad N, Pinsk V, Haim A, Fruchtman Y, et al. Abnormal neutrophil chemotactic activity in children with congenital insensitivity to pain with anhidrosis (CIPA): The role of nerve growth factor. Clin Immunol. 2009;

44. Indo Y, Mardy S, Tsuruta M, Karim MA, Matsuda I. Structure and organization of the human TRKA gene encoding a high affinity receptor for nerve growth factor. Jpn J Hum Genet. 1997;42(2):343–51.

45. Kaplan DR, Miller FD. Neurotrophin signal transduction in the nervous system. Curr Opin Neurobiol. 2000;

46. 1000 Genomes Project Consortium, Auton A, Brooks LD, Durbin RM, Garrison EP, Kang HM, et al. A global reference for human genetic variation. Nature. 2015;526(7571):68–74.

47. Lek M, Karczewski KJ, Minikel E V., Samocha KE, Banks E, Fennell T, et al. Analysis of protein-coding genetic variation in 60,706 humans. Nature. Nature Publishing Group; 2016;536(7616):285–91.

48. Walter K, Min JL, Huang J, Crooks L, Memari Y, McCarthy S, et al. The UK10K project identifies rare variants in health and disease. Nature. 2015;

49. Ogata H, Goto S, Sato K, Fujibuchi W, Bono H, Kanehisa M. KEGG: Kyoto encyclopedia of genes and genomes. Vol. 27, Nucleic Acids Research. 1999. p. 29–34.

50. Antonov A V, Schmidt EE, Dietmann S, Krestyaninova M, Hermjakob H. R spider: a network-based analysis of gene lists by combining signaling and metabolic pathways from Reactome and KEGG databases. Nucleic Acids Res. 2010 Jul 1;38(Web Server issue):W78–83.

51. Pereira R, Phillips C, Pinto N, Santos C, dos Santos SEB, Amorim A, et al. Straightforward inference of ancestry and admixture proportions through ancestry-informative insertion deletion multiplexing. PLoS One. 2012;7(1).

52. Zambrano AK, Gaviria A, Vela M, Cobos S, Leone PE, Gruezo C, et al. Ancestry characterization of Ecuador’s Highland mestizo population using autosomal AIM-INDELs. Forensic Sci Int Genet Suppl Ser. 2017;6.

53. RStudio Team. RStudio: Integrated Development for R. [Online] RStudio, Inc, Boston, MA URL http://www.rstudio.com. 2016;

54. Paz-Y-Miño C, López-Cortés A, Arévalo M, Sánchez ME. Monitoring of DNA damage in individuals exposed to petroleum hydrocarbons in Ecuador. Vol. 1140, Annals of the New York Academy of Sciences. 2008.

